## Supplemental Figures for "mTOR signaling regulates the morphology and migration of outer radial glia in developing human cortex"

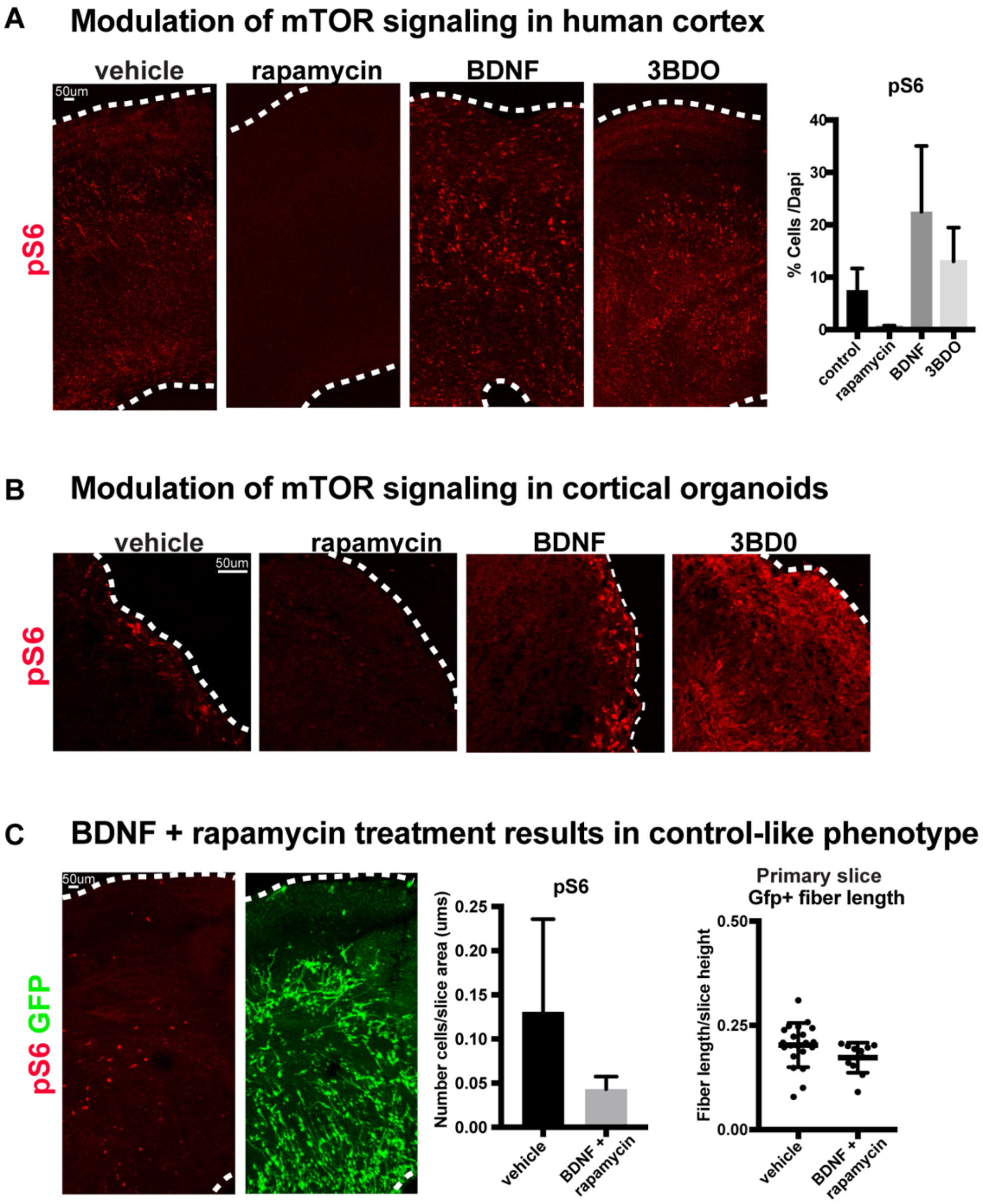


**Supplemental Figure 1: Manipulations of mTOR can be monitored through changes to pS6**
**A)** Primary human slice cultures express pS6 in the oSVZ. After rapamycin treatment there is little pS6 detected. After BDNF or 3BDO treatment, there is an increase in pS6+ cells throughout the culture (n>3 control, rapamycin, and BDNF and 3BDO-treated slices from three independent experiments; error bars represent SD). **B)** In organoids, pS6 is expressed at the edge of the organoid and rapamycin treatment results in complete loss of pS6. BDNF modestly increases pS6 levels, while 3BDO treatment increases pS6 expression throughout the organoid (n=6 control, rapamycin, BDNF and 3BDO-treated organoids from three PSC lines). **C)** When organotypic primary slice cultures are treated with BDNF and rapamycin together, pS6 expression returns to control-like levels. GFP+ fiber organization improves compared to either mTOR manipulation alone.

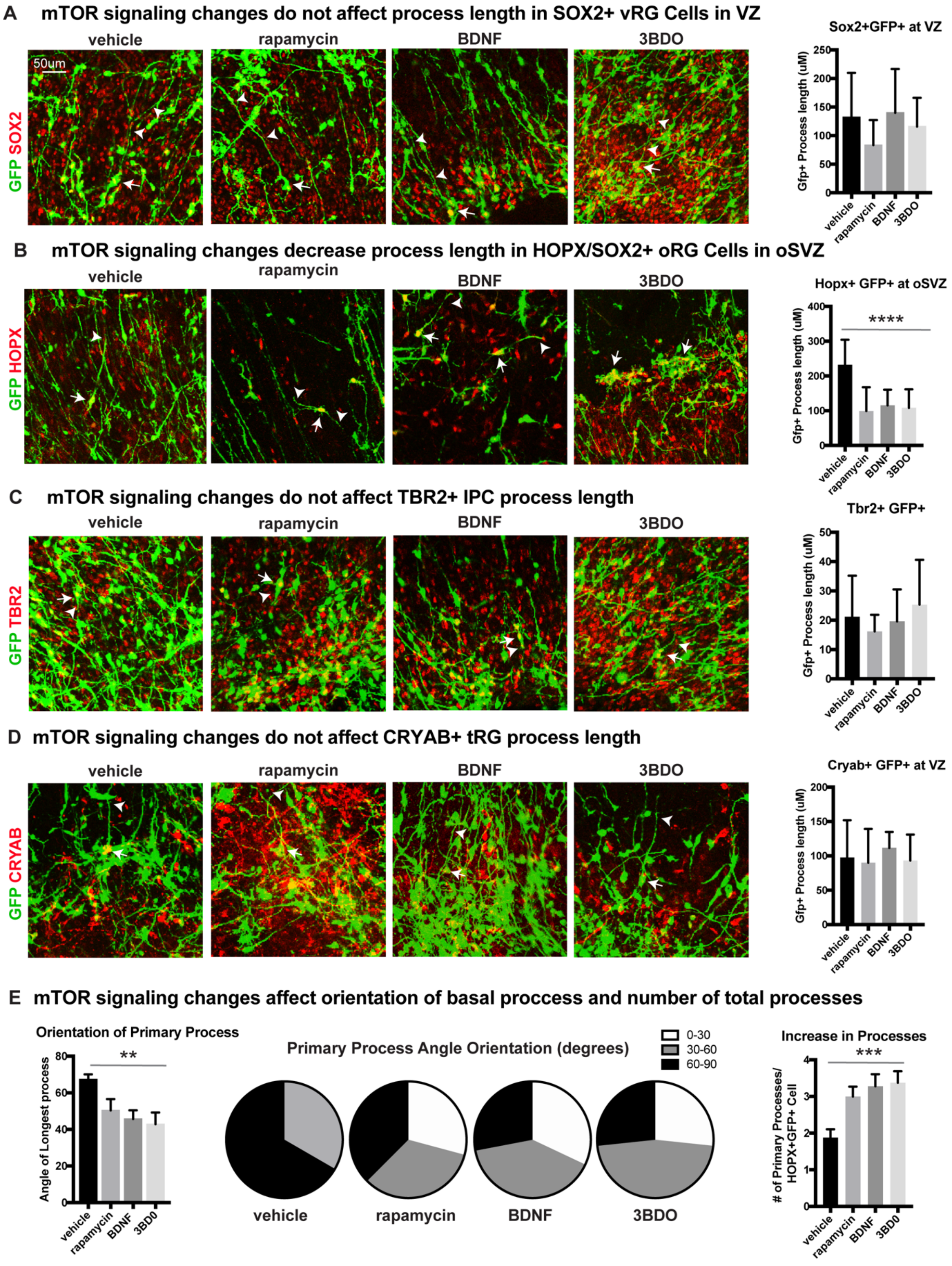


**Supplemental Figure 2: mTOR signaling has functional effects only on oRG progenitor cells**

**A)** mTOR signaling is manipulated using rapamycin, BDNF or 3BD0 in primary organotypic slice cultures infected with a CMV::GFP adenovirus to label the glial scaffold.There is no significant change to the GFP+ process length of SOX2+ GFP+ ventricular radial glial cells in the VZ (n>4 cells/slice across three independent experiments). **B)** There is a significant reduction of HOPX+GFP+ fiber length in oRG cells in the oSVZ in all mTOR treatment groups (n>4 cells/slice across three independent experiments; one-way ANOVA, ****p<0.0001). **C)** Manipulation of mTOR signaling does not affect GFP process length in TBR2+ GFP+ intermediate progenitor cells (n>4 cells/slice across three independent experiments). **D)** Changes to mTOR signaling do not affect CRYAB+ GFP+ fiber length of truncated radial glia (n>4 cells/slice across three independent experiments). **E)** Angle of primary process compared to the ventricular surface (0-90 degrees). mTOR manipulations affect the orientation of the basal fiber and decrease the average angle orientation (n>17cells/group across three independent experiments, one-way ANOVA,**p<0.01). Data also represented in proportions of cells from 0-30, 30-60, or 60-90 degrees per treatment group in pie charts. mTOR manipulations increase the number of processes (n>17 cells/group across three independent experiments, one-way ANOVA, ***p<0.001).

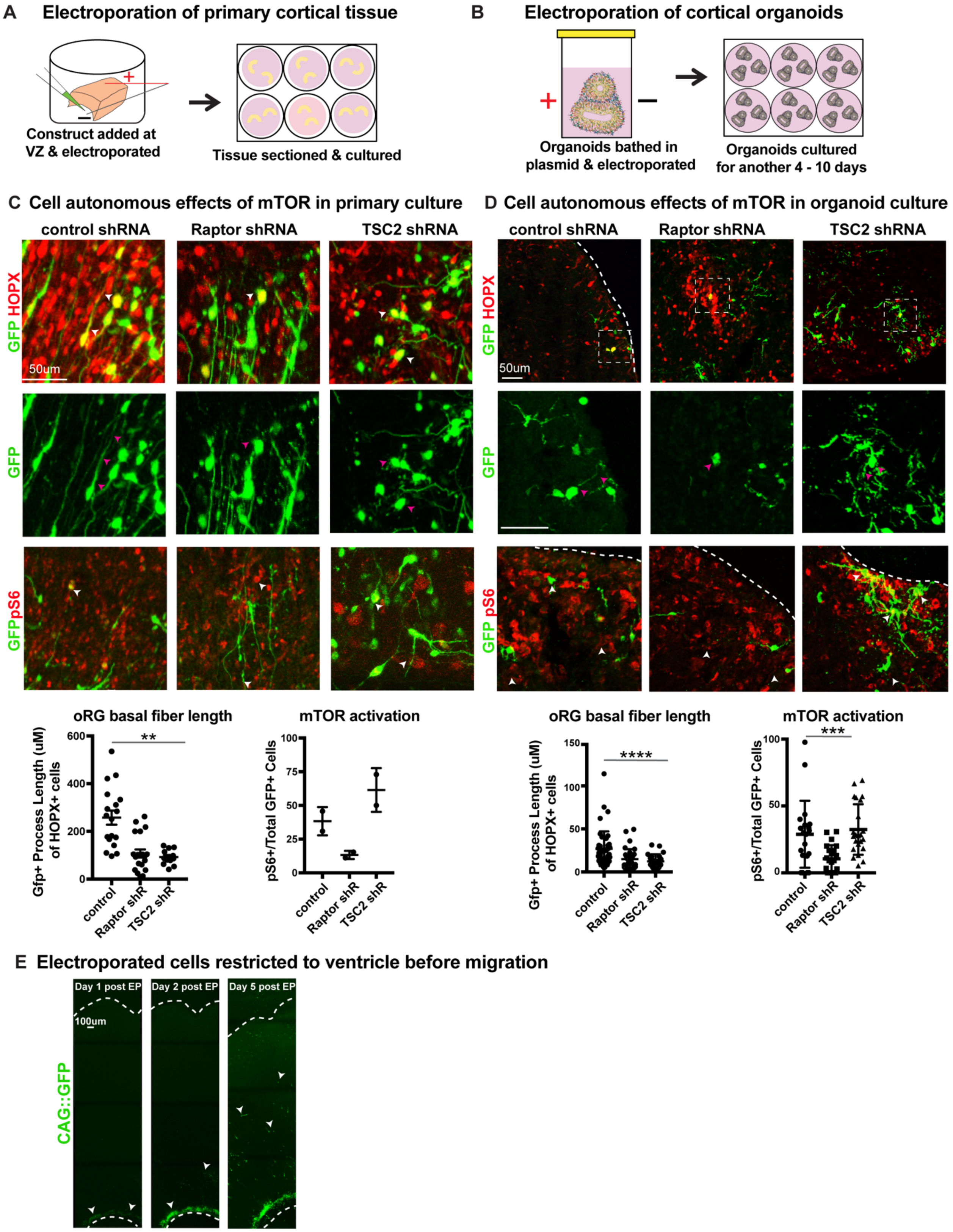


**Supplemental Figure 3: shRNAs against mTOR pathway components cell autonomously regulate oRG process length**

**A)** shRNAs were delivered along the ventricle of primary cortical tissue which was acutely sectioned and cultured for six days prior to collection. **B)** Organoids were bathed in media containing shRNAs,electroporated inside a cuvette and then cultured for 4 - 10 days prior to collection. **C)** Electroporation of either *RPTOR* or *TSC2* shRNAs resulted in a decrease of GFP+ process length in cells that were double-positive for HOPX (n=19 control, 20 Raptor and 12 TSC2-shRNA electroporated GFP+Hopx+ cells from three independent experiments; **$p<0.001$, $p=0.0532$, one-way ANOVA, error bars represent SEM). *RPTOR* electroporation resulted in a decrease in mTOR signaling indicated by pS6+, GFP+ cells, while TSC2 electroporation resulted in an increase in pS6+ GFP+ cells (n=2 tissue slices from two independent experiments; error bars represent SEM). **D)** Electroporation of *RPTOR* and *TSC2* shRNAs in cortical organoids. *RPTOR* and *TSC2* shRNA electroporation decreased the GFP+ primary process length in oRG cells. There were fewer pS6+ GFP+ cells after *RPTOR* shRNA electroporation, while *TSC2* electroporation had did not significantly affect the number of pS6+ GFP+ cells in the organoid (n>29 control, Raptor, and TSC2 electroporated GFP+HOPX+ cells from nine organoids per condition across four independent experiments; ****$p<0.0001$; ***$p<0.001$, one-way ANOVA, error bars represent SEM). **E)** Constructs are dropped along the ventricular edge of intact primary tissue and pulsed with electricity before sectioning and culturing. GFP+ electroporated cells are initially restricted to the ventricular edge demonstrated by their position one and two days post-electroporation. By five days post-electroporation GFP+ cells from the ventricular edge have migrated to the oSVZ (n>2 slices/group across two independent experiments).

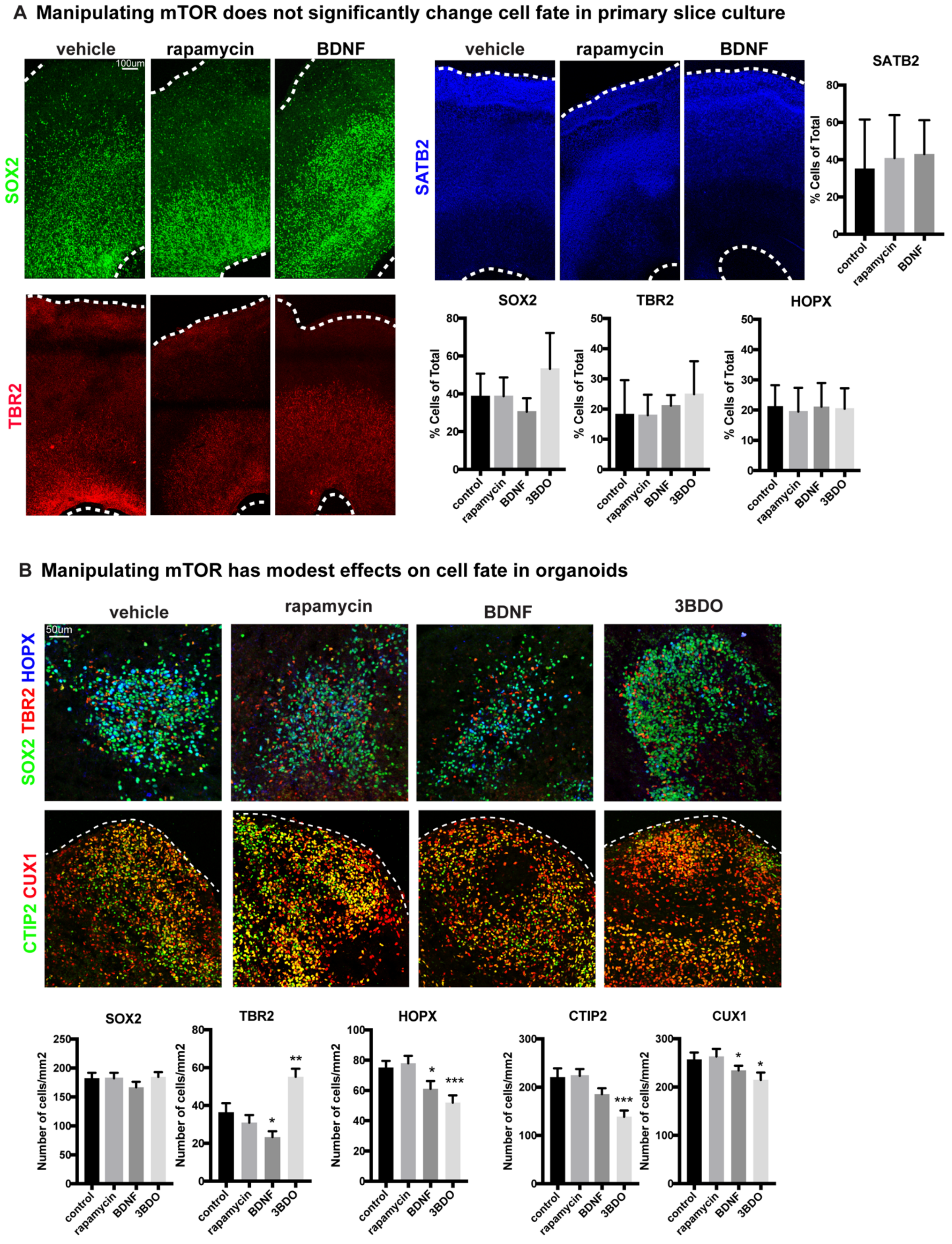


**Supplemental Figure 4: Manipulating mTOR signaling has little effect on cell fate**

**A)** Activating or inhibiting mTOR in primary slice cultures has no effect on cell fate. There is no significant change in SOX2+ progenitors, HOPX+ oRGs, TBR2+ IPCs or SATB2+ upper layer layer neurons (n>3slices/group/stain, Sox2: $p=0.15$, Tbr2: $p=0.62$, Hopx: $p=0.95$, Satb2: $p=0.91$ using one-way ANOVA, error bars represent SD). **B)** Changes to mTOR signaling have modest effects on cell fate in organoids. There is no change to SOX2+ cells in any condition and rapamycin has no effect on any of the cell types assessed. mTOR activators, BDNF and 3BDO, have modest effects decreasing HOPX+ oRGs, CTIP2+ lower layer neurons, and CUX1+ upper layer neurons. BDNF modestly decreases TBR2+ IPCs, while 3BDO increases them. The number of cells positive for each marker was counted and then normalized to organoid size (n>50sections/group from six organoids across three lines for all stains; *$p<0.05$, **$p<0.01$, ***$p<0.001$; student's two-tailed t-test per group compared to control, error bars represent SEM).

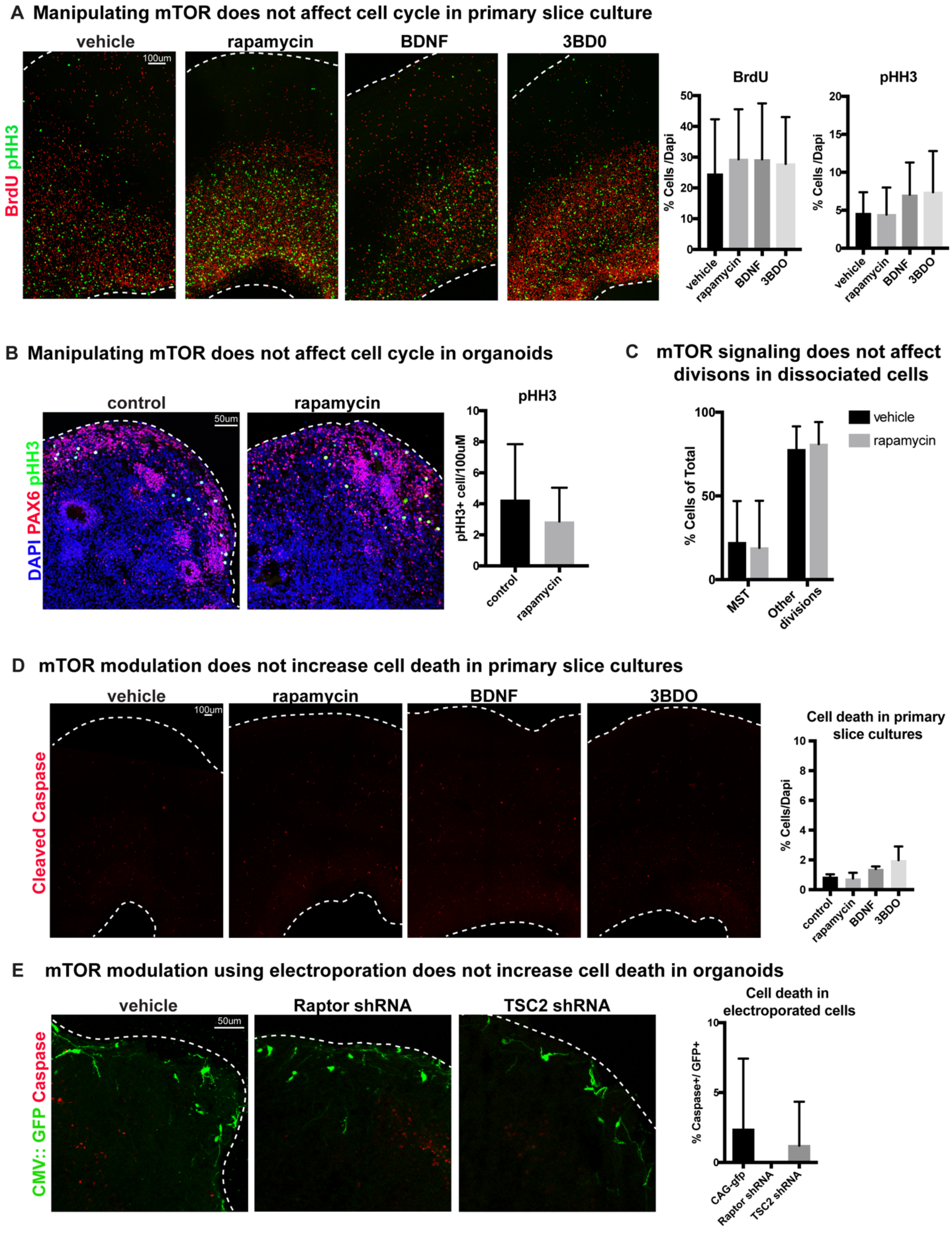


**Supplemental Figure 5: Changes to mTOR signaling do not affect cell cycle or cell death**

**A)** Manipulating mTOR signaling has no effect on BrdU+ progenitor cells in synthesis or pHH3+ mitotic cells in human slice cultures (n>3 slices/group across five independent experiments; BrdU: p=0.98, pHH3: p=0.54, one-way ANOVA, error bars represent SD). **B)** Inhibiting mTOR signaling has no effect on the number of pHH3+ cells in mitosis in organoids. Counts were normalized to organoid size (n>19 sections/group from four organoids; p=0.12, error bars represent SD). **C)** Rapamycin treatment has no effect on the ability of cells to divide and does not change the frequency of oRG MST in dynamically imaged primary radial glia (p=0.99, two-tailed student's t-test, error bars SD). **D)** Manipulating mTOR signaling does not increase the prevalence of cleaved caspase-3 in any condition (n=3 slices/group, p=0.2, one-way ANOVA, error bars represent SD). **E)** Electroporation of mTOR shRNA constructs does not increase the number of GFP+ cleaved caspase+ cells compared to control electroporations (n=4 organoids/group, p=0.44, one-way ANOVA, error bars represent SD).

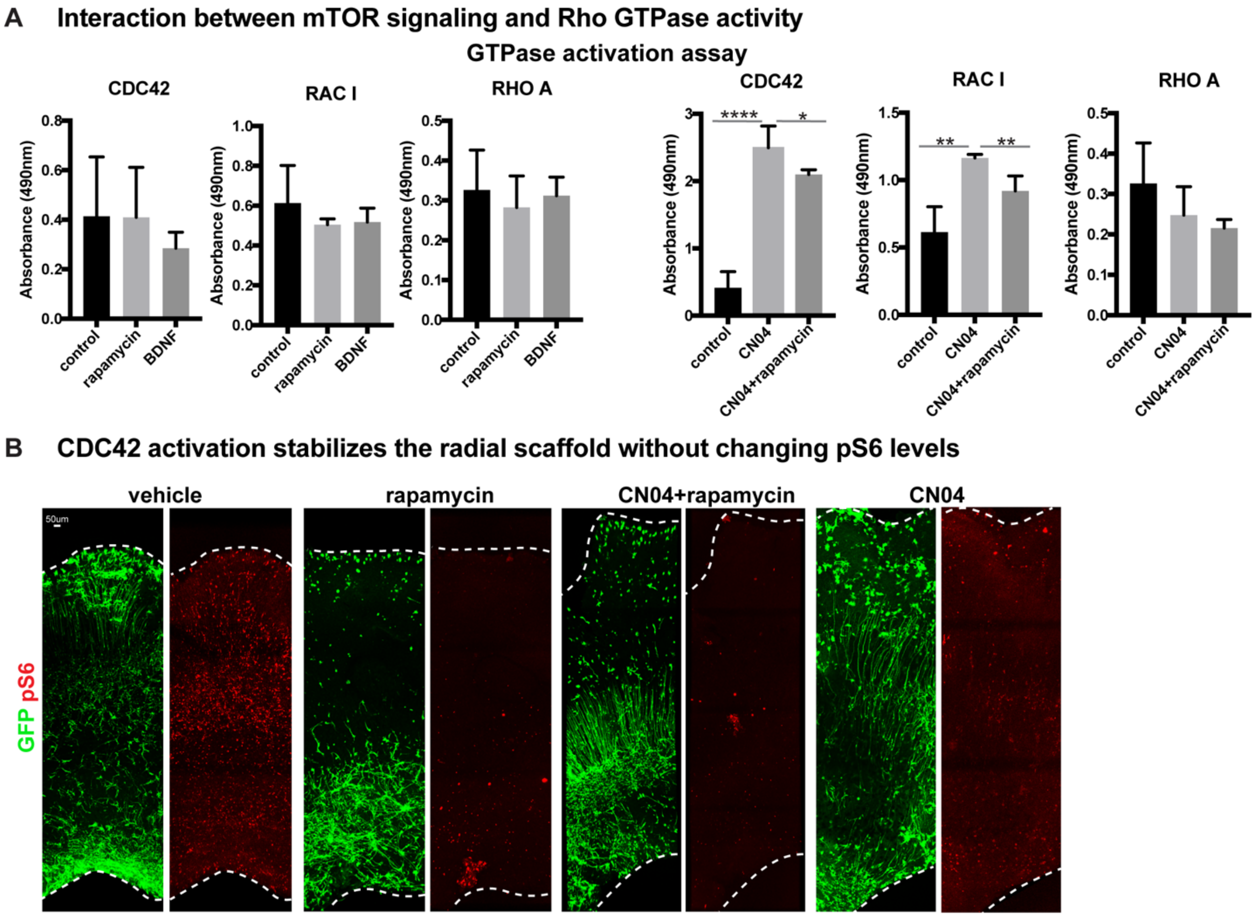


**Supplemental Figure 6: mTOR signaling modulation changes CDC42 and RAC1 activity**

**A)** The VZ to oSZV of primary slice cultures, treated with rapamycin, BDNF, CN04 or CN04+rapamycin, were microdissected, isolated and protein extracted. Protein GTPase activation assays demonstrate little baseline change to any Rho GTPase after mTOR manipulation. CN04 activates both CDC42 and RAC1, and treatment of CN04 in combination with rapamycin results in an intermediate level of activity showing interaction between the two pathways (three independent experiments/protein/group, *$p<0.05$, **$p<0.01$, ****$p<0.0001$, unpaired two-tailed student's t-tests were performed to compare each group to the control). **B)** Inhibiting mTOR signaling results in a reduction of downstream pS6 activity. Although Cdc42 activity rescues mTOR-mediated oRG morphology, CN04 does not rescue the loss of pS6 in combination with rapamycin nor does CN04 alone change pS6 levels (three slices/group across three independent experiments).
